# The large conjugative plasmid pAB3 integrates into the chromosome of *Acinetobacter baumannii* ATCC 17978

**DOI:** 10.64898/2026.10.30.763009

**Authors:** Matthew Neil, Gemma C. Langridge, Benjamin A. Evans

## Abstract

ATCC 17978 is one of the most utilised lab strains of *Acinetobacter baumannii*. Current literature suggests ATCC 17978 can have either a sulfamethoxazole-resistant phenotype or have the type VI secretion system (T6SS) switched on depending on the presence/absence of the large conjugative plasmid pAB3, which encodes the antimicrobial resistance gene (ARG) *sul2* and T6SS repressors *tetR1*/*tetR2*. Here, we show that pAB3 integrates into the ATCC 17978 chromosome, which enables a large inversion resulting in the chromosomal arrangement labelled GS2.24 by Socru. This integration has no transcriptional impact on nearby genes or *tetR1*/*tetR2*. We find several instances were a portion of integrated pAB3 containing *tetR2* is lost but the rest of the sequence, including *sul2*, is maintained, suggesting pAB3 integration may serve to make further rearrangements more tractable, enabling a putative T6SS+ sulfamethoxazole-resistant phenotype.

**Impact statement:** *Acinetobacter baumannii* is a major cause of hospital infections and is often resistant to antibiotics. The strain ATCC 17978 is one of the most widely used laboratory strains, underpinning research on how this bacterium competes with other microbes, causes disease and resists treatment. Its original genome assembly placed a plasmid, pAB3, within the chromosome, which was later considered an error. We show that pAB3 is genuinely integrated into the chromosome in just over half of the ATCC 17978 strains studied. This integration enables a large chromosomal inversion but does not alter the activity of nearby genes under standard conditions. We also identify strains in which a region of pAB3 encoding a repressor of the type VI secretion system, a bacterial weapon, is missing while a sulfamethoxazole resistance gene is retained, with implications for trade-offs between bacterial virulence and antimicrobial resistance. These findings reveal that the ATCC 17978 chromosome is more variable than previously recognised, a significant advance for researchers using this strain, and contributes to understanding of bacterial chromosome structural variation, plasmid integration and resistance evolution, of interest to microbiologists, clinicians and evolutionary biologists.

**Data summary:** Sequencing data is available on the sequence read archive (SRA) under BioProject PRJNA1544396.

## Introduction

*Acinetobacter baumannii* is a highly antimicrobial resistant (AMR) pathogen which is a major cause of nosocomial infections (1). A central determinant of the pathogenicity of *A. baumannii* is the type VI secretion system (T6SS), which primarily acts to kill competing bacteria, but has also been linked to virulence and AMR through its role in modulating horizontal gene transfer (HGT) (2). Thus, *A. baumannii* isolates can be labelled as T6SS+ and T6SS-to differentiate isolates with the T6SS on (T6SS+) or off (T6SS-) respectively (3,4). Expression of the T6SS is repressed by TetR1 and TetR2, located on the large conjugative plasmid pAB3, which also encodes the antimicrobial resistance gene (ARG) *sul2* which confers a sulfamethoxazole-resistant phenotype (3), and contains a sequence derived from a common ancestor with the AbGRI1 chromosomally-located resistance island (5). Spontaneous loss of pAB3 (which has been documented as occurring during transport between laboratories (6)) enables a switch from a sulfamethoxazole-resistant T6SS- (T6SS off) phenotype to a non-sulfamethoxazole-resistant T6SS+ (T6SS on) phenotype (3).

One of the most common strains of *A. baumannii* studied *in vitro* is ATCC 17978 (7), which belongs to multi locus sequence typing (MLST) sequence type (ST) 437 (8). Several genomic variants of ATCC 17978 are known to exist which have been reviewed elsewhere (7). Genomic variants in bacteria range from small-scale single nucleotide polymorphisms (SNPs) to large-scale structural variants (SVs), the latter of which are less frequently discussed but known to be common in bacterial chromosomes (9,10). Many SVs arise from inversions around the origin of replication (oriC) (10) mediated by homologous recombination around inverted repeat (IR) sequences (11). The bias towards these inversions results from selection for a chromosome to have equally sized replichores i.e. with the origin and terminus of replication being 50% of the chromosome length apart, usually termed “balanced”. Chromosomal imbalance can be tolerated, though typically extremely imbalanced chromosomes incur a fitness cost (12,13).

In the original ATCC 17978 genome (14) pAB3 was assembled as part of the chromosome, although this has since been deemed a misassembly in the literature (3,5–7,15,16). Here, we show that pAB3 is integrated into the ATCC 17978 chromosome in just over half of strains studied. The integration of pAB3 supplies an IS*Aba18* element which enables a large inversion to occur approximately symmetrically around the oriC. However, this integration does not transcriptionally impact nearby genes or *tetR1*/*tetR2* under baseline conditions. We find several instances where a section of the integrated pAB3 containing *tetR2* is absent, suggesting that the integration of pAB3 may serve to enable further structural variation in order to de-repress the T6SS while maintaining part of the pAB3 sequence encoding *sul2*. This would enable a putative T6SS+ sulfamethoxazole-resistant phenotype. Overall, we show that the ATCC 17978 chromosome is significantly more variable than previously recognised.

## Methods

### Dataset curation

A set of *A. baumannii* chromosomes curated and annotated as part of a previous study (17) were filtered to include long-read or hybrid sequenced, *de novo* assembled genomes only. ST437 chromosomes were considered ATCC 17978 isolates.

### Socru

To determine the structural arrangement of each chromosome, Socru v2.25 (9) was run on all chromosomes with species set to “Acinetobacter_baumannii” and all other parameters kept as default.

### Chromosome comparison

Alignments between chromosomes were obtained with MEGABLAST implemented in BLAST v2.17.0 (18) with the E-value threshold set at 1e-10. Alignments under 500 bp in length were removed. Alignments were visualised with pyGenomeViz v1.6.1 (https://github.com/moshi4/pyGenomeViz) and pyCirclize v1.10.1 (https://github.com/moshi4/pyCirclize).

### Breakpoint identity detection

In all ST437 GS2.24 isolates, a 40 kbp sequence composed of 20 kbp upstream and 20 kbp downstream was extracted from both breakpoints in each chromosome. One sequence was reverse complemented and aligned to the other using the EMBOSS water aligner implemented in EMBOSS v6.6.0 (19). From this alignment, the longest run of identical nucleotides was extracted and its identity was determined with the TnCentral database (20).

### pAB1, pAB2, and pAB3 detection

Initial pAB3 detection was performed by extracting the DNA sequence of the insertion into the region defined by Socru as fragment 6 (Fig. 1) and querying against the NCBI core nucleotide database (21). To more accurately determine the locations of plasmid integration and contig identity, the sequences of pAB1, pAB2, and pAB3 (GenBank accessions CP079933.1, CP079934.1 CP079932.1 respectively) were aligned against publicly available and hybrid assembled ST437 genomes with an E-value threshold of 1e-10 and an identity threshold of 99%. To avoid spurious results, hits under 500 bp long were removed, and hits within 200 bp of each other were merged into blocks. To avoid erroneously detecting AbGRI1 as pAB3, blocks under 30 kbp in length were excluded. Contigs where a single block covered 95% or more of both the contig and plasmid were considered to be that plasmid.

**Fig. 1.**
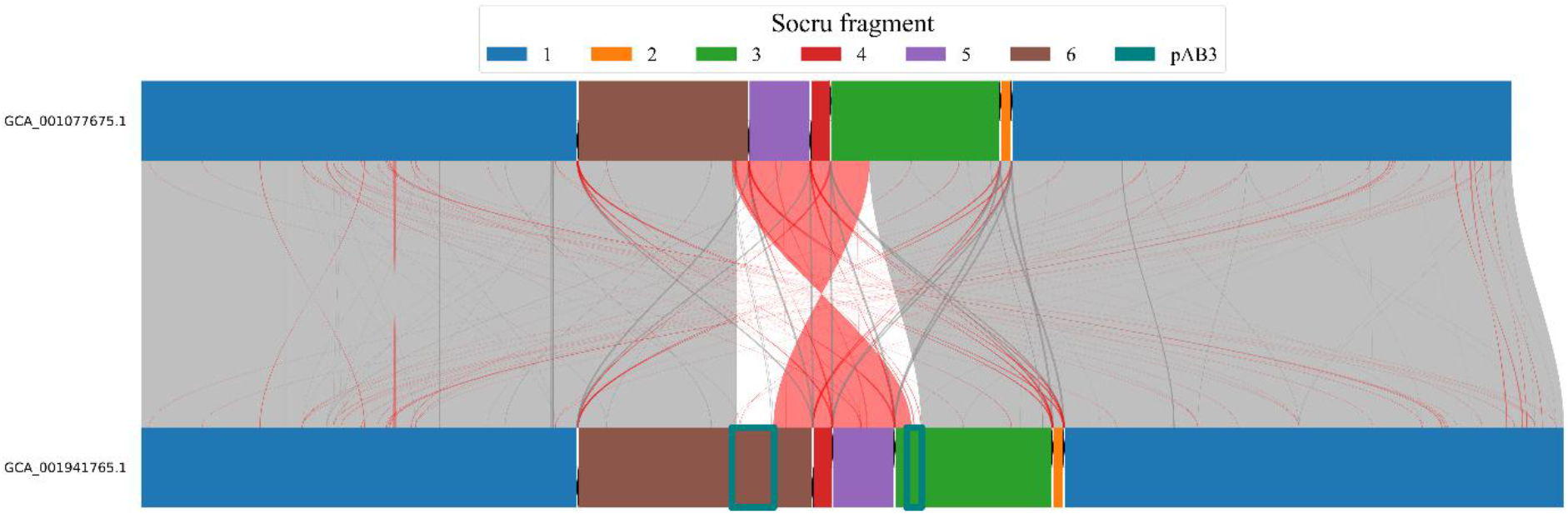
Comparison between GS1.0 (top) and GS2.24 (bottom) chromosomes. Aligned regions <500 bp are not shown. Standard alignments are shown with grey links, while inverted alignments are shown with red links. rRNA operons are shown with black arrows between fragments indicating the direction determined by Socru, and fragments are colour coded as per the legend. The *dnaA* gene is shown in magenta, located in fragment 4. pAB3 sequences are shown with teal boxes.

**Fig. 2.**
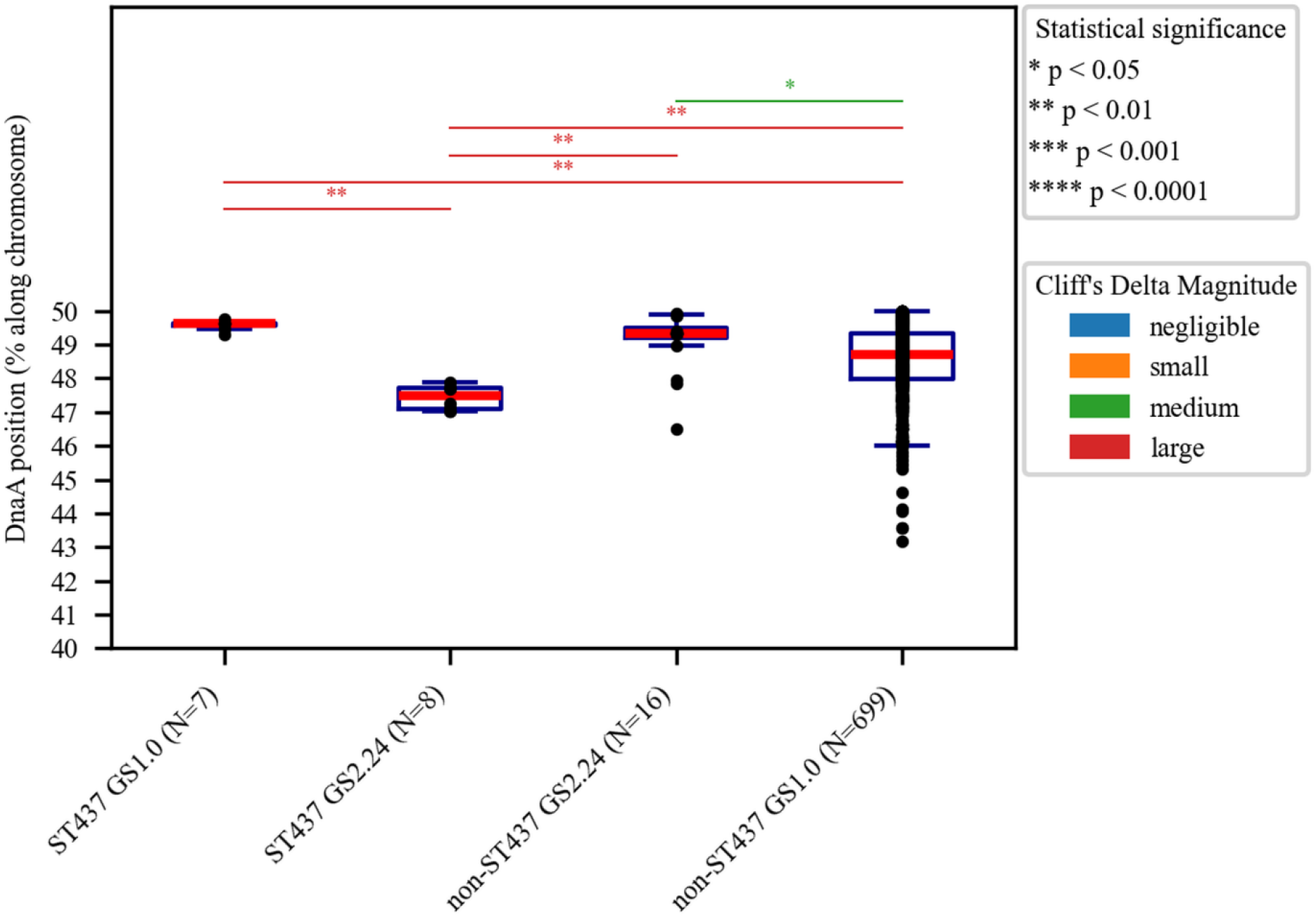
Balance of ST437/non-ST437 G1.0/GS2.24 chromosomes, as measured by *dnaA* percentage distance from ter. Individual data points are shown with black circles. Each Boxplot shows the median (red line), interquartile range (IQR) (box), and whiskers extending to 1.5×IQR from the quartiles. Statistical significance (p<0.05) was determined through the Mann-Whitney U test with the Holm-Bonferroni correction applied. Effect size was determined through Cliff’s delta.

### *tetR1* and *tetR2* detection

Protein sequences for TetR1 and TetR2 were extracted from Protein Data Bank (PDB) structures (8Z6D and 8Z6E respectively), created as part of a previous study (22). TBLASTN, implemented in BLAST as above, was used to locate *tetR1* and *tetR2* genes on DNA sequences with an identity threshold of 90% and a coverage threshold of 95%. Query sequences starting more than 2 bp from the reference sequence start were excluded.

### *comM* detection *in silico*

Forward and reverse primer sequences targeting the 5’ and 3’ portions of the *comM* locus were obtained from a previous study (23). The locations of these probes were determined with BLASTN, implemented in BLAST as described above, with BLASTN-short default parameters. A bitscore threshold of 30 and an E-value threshold of 0.01 was set. For all chromosomes the 5’ and 3’ primer pairs (forward and reverse) were located very close together, so the midpoints between each pair were annotated as comM_1 and comM_2 respectively.

### Statistical testing

Mann-Whitney U tests were performed using Scipy v1.17.1 to determine whether differences in chromosomal balance were statistically significant between different subsets of *A. baumannii*. Effect sizes were determined with Cliff’s delta implemented in cliffsDelta v1.0.0 (https://github.com/neilernst/cliffsDelta). *p* values were adjusted for multiple testing with the Holm-Bonferroni correction implemented in statsmodels v0.14.6 (24).

### Isolate culturing and DNA extraction

A strain corresponding to the accession GCA_001077675.1 (ATCC 17978-mff) was kindly provided by Mario Feldman, while strains corresponding to accessions GCA_001941765.1 (AB042), and GCA_001593425.2 (ATCC 17978-VU) were kindly provided by Ayush Kumar. The minimum inhibitory concentration (MIC) of meropenem was determined for each strain by measuring growth in Mueller Hinton Broth across a range of concentrations via OD_600_ measurement according to EUCAST guidelines. MIC was defined as the minimum concentration of meropenem which completely inhibited growth. Strains were then cultured in either no antibiotic or in a sub-MIC of meropenem (17978-mff, 0.06 mg/L; AB042, 0.5 mg/L; 17978-VU, 0.25 mg/L). Genomic DNA was extracted using the Wizard Genomic DNA

Purification Kit (Promega, Madison, WI, USA) following the manufacturer’s protocol for Gram-negative bacteria with the minor modification that protein precipitation solution was ice-cold and added to extraction tubes on ice.

### Long read whole genome sequencing

200 ng of DNA per sample was repaired using the NEBNext FFPE DNA Repair Mix and end-prepped using the NEBNext Ultra II End Prep modules, then barcoded using the Oxford Nanopore Native Barcoding Kit 96 V14 (SQK-NBD114.96) with NEB Blunt/TA Ligase Master Mix. Barcoded libraries were pooled, cleaned with 0.6X SPRI beads, ligated to sequencing adapters using the NEBNext Quick Ligation Module, cleaned again with 0.4X SPRI beads and Short Fragment Buffer, and sequenced on an R10.4.1 flow cell (FLO-PRO114M) on a PromethION 2 Solo. Reads were basecalled using the R10.4.1 super-accuracy model (dna_r10.4.1_e8.2_400bps_sup v5.0). Adaptors were trimmed with porechop ABI v0.5.1 (25) and initial quality control metrics were obtained with seqkit v2.10.0 (26).

### Short read whole genome sequencing

Genomic DNA was normalised to 5 ng/µl. Libraries were prepared using a miniaturised Illumina DNA Prep protocol. Briefly, 10 ng of DNA was tagmentated with Bead Linked Transposomes and amplified for 14 cycles using KAPA 2G Fast Hot Start Ready Mix and Illumina IDT unique dual index primers. Libraries were quantified using the QuantiFluor dsDNA System, pooled equimolarly, and size-selected using a double-sided 0.5-0.7X SPRI clean-up. The final pool was quantified by Qubit and Agilent TapeStation (D5000 ScreenTape) and sequenced on the Illumina NextSeq 2000 platform at a final concentration of 750 pM, generating 2×151bp paired-end reads. Basecalling and adaptor trimming was conducted with DRAGEN BCL Convert v4.2.7 and initial quality control metrics were obtained with seqkit v2.10.0 (26).

### Hybrid genome assembly and quality control

Long reads were filtered to remove reads <1 kbp long and/or <Q20 using chopper v0.13.0 (27). Reads were then downsampled to an estimated 80X coverage prioritising read length over quality with Filtlong v0.3.1 (https://github.com/bongadi/Filtlong), with the corresponding publicly available assembly used to estimate genome length. Short reads were filtered to remove reads <50 bp and/or <Q20 using chopper as above. Hybrid assembly was performed using Unicycler v0.5.1 (28) in bold, normal, and conservative modes. Average nucleotide identity (ANI) to an *A. baumannii* type strain (ATCC 19606) genome (GenBank accession: CP045110.1) was determined with FastANI v1.34 (29). Benchmarking universal single-copy orthologs (BUSCO) scores for both the longest contig only and all contigs were obtained using BUSCO v6.1.0 (30) using the “acinetobacter_odb12.2” lineage dataset.

### Hybrid chromosome reorientation, annotation, and comparison

For each isolate, the longest contig from the conservative Unicycler run was extracted and considered to be the chromosome. Chromosomal dif site sequences produced as part of a previous study (31) were used to generate a position-specific scoring matrix (PSSM) with Biopython v1.87 (32), which was in turn used to reorient each chromosome to start at the dif site. The *dnaA* gene was identified with TBLASTN, implemented in BLAST as above, and chromosomes were reverse complemented as needed such that the *dnaA* gene was on the positive strand. Chromosomes were annotated using bakta v1.12.1 (33) with the heavy database and digIS v1.2 (34). Multi-locus sequence typing (MLST) was performed with FastMLST v0.0.19 (35) according to the Pasteur scheme. ARGs were detected with the resistance gene identifier (RGI) v6.0.8 which queries the comprehensive antibiotic resistance database (CARD) v4.0.2 (36). NucDiff v2.0.3 (37) was used to detect both SNP and SV differences between all public and hybrid assembled chromosomes.

### RNA sequencing, trimming, and filtering

RNA extraction and library preparation was performed by Lexogen following their standard protocol. Libraries were indexed using Lexogen unique dual indices (UDIs) and sequenced on a NovaSeq X (Illumina) using a 5B flow cell in SR100 mode. Reads were basecalled using DRAGEN-based real-time analysis (RTA4). Adaptor trimming was performed according to the CORALL analysis pipeline (https://github.com/Lexogen-Tools/corall_analysis). During this stage it was found that the high number of reads aligning to the transfer-messenger RNA (tmRNA) of each genome was resulting in unsustainably high memory usage, so this region was masked in all samples, leading to the tmRNA being excluded from the final data. Reads <50bp and/or <Q20 were discarded with chopper as above.

### Differential gene expression analysis

RNAseq reads were aligned to their respective hybrid genome using BWA-MEM v0.7.19-r1273 (38), and the alignments were sorted and indexed using samtools v1.24 (39). featureCounts (40), implemented as part of subread v2.1.1, was run using a bakta-produced annotation with feature type “CDS” and attribute type “locus_tag”. Strand-specific counting (- s 1) was used as the protocol produces reads in the same orientation as the original RNA. The resulting raw counts were fed into DESeq2 v1.50.2 (41), implemented in R v4.5.3, and a significance threshold of 0.05 was applied to identity differentially expressed genes (DEGs).

## Results

### ATCC 17978 chromosomes have two structural arrangements

An initial dataset of 1,095 publicly available *A. baumannii* chromosomes was produced by filtering a previously constructed dataset (17) to only long-read or hybrid sequenced *de novo* assembled genomes. Of these, 15 were ST437, considered strains of ATCC 17978 (8). Of these, 13 are described as ATCC 17978 strains, and two are known derivatives of an ATCC 17978 strain (Table 1). After running Socru on these chromosomes it was found that 7 are in the standard arrangement (GS1.0) and 8 are in an arrangement corresponding to GS2.24. Alignment of a GS1.0 reference (GenBank accession: GCA_001077675.1, strain: ATCC 17978-mff) against a representative GS2.24 (GenBank accession: GCA_001941765.1, strain: AB042) showed that the GS2.24 arrangement differs by a large inversion centered approximately on oriC, with breakpoints in fragments 3 and 6, and two insertions in fragments 3 and 6 approximately 50 kbp and 100 kbp long respectively (Fig. 1).

**Table 1.** Summary of ST437 chromosomal arrangement, pAB3 integration status, *tetR1*/*tetR2* presence/absence, and chromosomal IS*Aba18* copy number. “GS id” refers to the Socru id for the chromosome structural arrangement. “None” specifies no pAB1, pAB2, or pAB3 plasmids were identified in the isolate genome.

| Accession | Strain | GS id | plasmids | pAB3 integrated into chromosome? | *tetR1* | *tetR2* | IS*Aba18* copy number |
|---|---|---|---|---|---|---|---|
| GCA_004794235.2 | ATCC 17978 | GS2.24 | pAB1, pAB2 | yes | yes | no | 2 |
| GCA_013372085.1 | ATCC 17978 | GS2.24 | pAB1, pAB2 | yes | yes | yes | 2 |
| GCA_029459615.1 | ATCC17978 TnAraC-H0N27_12600 | GS2.24 | pAB1, pAB2 | yes | yes | yes | 2 |
| GCA_004797155.2 | ATCC 17978 | GS2.24 | pAB2 | yes | yes | no | 2 |
| GCA_014672775.1 | ATCC 17978 | GS2.24 | pAB2 | yes | yes | yes | 2 |
| GCA_001941765.1 | AB042 | GS2.24 | None | yes | yes | yes | 2 |
| GCA_004794275.2 | ATCC 17978 | GS2.24 | None | yes | yes | yes | 2 |
| GCA_008630895.1 | AB043 | GS2.24 | None | yes | yes | yes | 2 |
| GCA_019356215.1 | 17978UN | GS1.0 | pAB1, pAB2, pAB3 | no | yes | yes | 0 |
| GCA_020912005.1 | ATCC 17978 | GS1.0 | pAB2, pAB3 | no | yes | yes | 0 |
| GCA_001077675.1 | ATCC 17978-mff | GS1.0 | pAB3 | no | yes | yes | 1 |
| GCA_026167785.1 | ATCC 17978 | GS1.0 | pAB3 | no | yes | yes | 0 |
| GCA_026167805.1 | ATCC 17978 | GS1.0 | pAB3 | no | yes | yes | 0 |
| GCA_019316825.1 | Ab17978UN | GS1.0 | None | no | N/A | N/A | 0 |
| GCA_022467135.1 | ATCC17978-VUB | GS1.0 | None | no | N/A | N/A | 0 |

### The large conjugative plasmid pAB3 is integrated into the chromosome of GS2.24 ATCC 17978 isolates

Querying the sequence of the DNA inserted into fragment 6 against the NCBI database gave hits back to the large conjugative plasmid pAB3. Further analysis of the 100 kbp block from each chromosome found instances where a segment containing *tetR2* was missing (Table 1). To determine the relationship between pAB3 integration, plasmid content and chromosomal arrangement, the pAB1, pAB2, and pAB3 plasmid sequences were aligned against the full genome of all 15 ST437 isolates in the dataset. The results are shown in Table 1. All ST437 genomes where pAB3 was integrated into the chromosome did not have a separate pAB3 contig and had the GS2.24 structural arrangement.

### The GS2.24 inversion breakpoint is IS*Aba18*

Further inspection of the alignments between GS1.0 and GS2.24 chromosomes showed that the inversion breakpoints are not at the rRNA operons as expected based on previous literature (9). To identify the genomic feature acting as an IR enabling this inversion, flanking sequences around both the fragment 6 and fragment 3 breakpoints were extracted. One sequence was reverse complemented and aligned against the other (see Methods), with the longest run of fully conserved bases being identified as IS*Aba18* in all cases. Therefore, it is highly likely that IS*Aba18* acted as the IR to enable the inversion leading to the GS2.24 arrangement. This is supported by the fact that GS2.24 ST437 isolates consistently have 2 copies of IS*Aba18*, while GS1.0 ST437 isolates have 0 or 1.

### pAB3 integration does not make chromosomes more balanced

Bacterial chromosomes typically have equal sized replichores (termed “balanced”) with the oriC ∼50% of chromosome length away from the ter (see Introduction). It was initially hypothesised that the integration of pAB3 may serve to balance the chromosome, as the GS2.24 inversion involves the movement of fragment 4 (containing the oriC) and fragment 5. Non-ST437 GS2.24 isolates were found not to have pAB3 integrated into the chromosome, with many having GS2.24 inversion breakpoints inside fragment 3 and fragment 6 (i.e. not at rRNA operons). Therefore, these were used to test this hypothesis. ST437 GS2.24 chromosomes had lower median *dnaA* distance from ter than non-ST437 GS2.24 chromosomes, showing that pAB3 integration and GS2.24 inversion actually makes the chromosome less balanced. An *in silico* generated, hypothetical mff chromosome (GS1.0) with pAB3 artificially integrated but no GS2.24 inversion was found to be more balanced (∼48.5%) than ST437 chromosomes with the GS2.24 inversion (∼47.5%), demonstrating that if the GS2.24 inversion occurred after pAB3 insertion this would not rebalance the chromosome. Overall, this shows that the integration of pAB3 and the GS2.24 inversion both make the chromosome more imbalanced. It should be noted that the replichores are still close to equal size in these chromosomes, and the *dnaA* positions are within the range found for non-ST437 G1.0 chromosomes. Therefore, it is likely that this apparent unbalancing does not affect fitness.

### Hybrid assemblies confirm the integration of pAB3 into the ST437 chromosome

As the SVs described above were based on publicly available genomes, the possibility that they were misassemblies arising by reference-guided genome assembly using the original ATCC 17978 genome (considered to be misassembled in current literature) could not be ruled out. Therefore, three isolates (GCA_001077675.1, strain: ATCC 17978-mff; GCA_001941765.1, strain: AB042; GCA_001593425.2, strain: ATCC 17978-VU) were selected for sequencing. ATCC 17978-mff, referred to herein as “mff”, is a commonly used reference strain (6,42) with a GS1.0 structure. ATCC 17978-VU, referred to herein as “VU”, was not part of our original dataset as its genome was not long-read sequenced, but was selected as it shows a GS2.24 structure. AB042 is a triclosan-resistant mutant (43,44) of ATCC 17978-VU.

All three isolates were long-read and short-read sequenced, then *de novo* assembled with Unicycler on bold, normal, and conservative modes. Chromosome and pAB3 contig lengths were the same for all three modes, with the main difference being that the pAB1 and pAB2 plasmids were fused together into a single contig on bold mode in mff and AB042. In the VU strain this was observed on all three modes. The conservative assemblies for all three isolates were carried forward for further testing. ANI to an *A. baumannii* ATCC 19606 strain genome was >95% for all genomes. BUSCO scores were >99% for all genomes and were identical to the BUSCO scores for chromosomes on their own. Overall, this shows high-quality chromosomes were assembled.

Testing the hybrid genomes for pAB3 integration revealed similar results to the publicly available genomes for mff and AB042, namely that in mff (GS1.0) pAB3 is found as a separate ∼150 kbp contig representing the independent plasmid, while in AB042 (GS2.24) pAB3 is integrated into the chromosome as two blocks of 100 kbp and 50 kbp. The VU chromosome contained a 50 kbp pAB3 block, but the 100 kbp block was missing (Fig. 3). This resulted in the *tetR1* and *tetR2* genes being lost. Altogether this showed that the publicly available chromosomes for these strains do not represent misassemblies.

**Fig 3.**
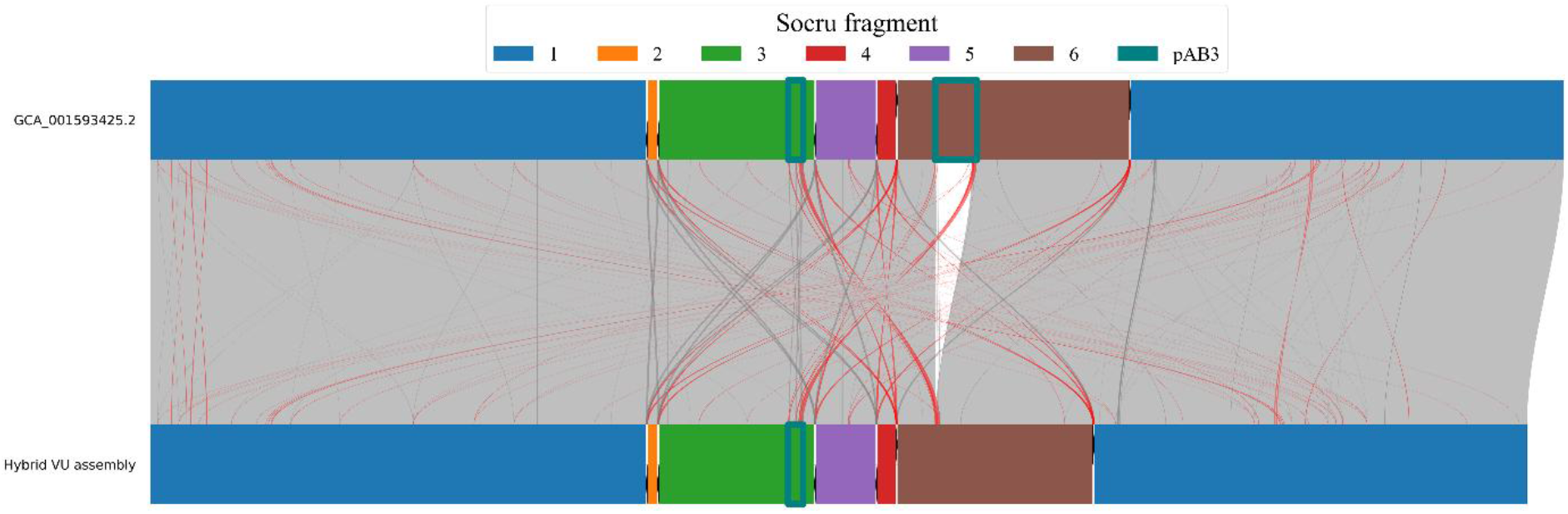
Comparison between publicly available (top) and hybrid assembled VU (bottom) chromosomes. Aligned regions <500 bp are not shown. Standard alignments are shown with grey links, while inverted alignments are shown with red links. rRNA operons are shown with black arrows indicating the direction determined by Socru, and fragments are colour coded as per the legend. The *dnaA* gene is shown in magenta in fragment 4. pAB3 sequences are shown with teal boxes.

### Integration of pAB3 does not transcriptionally impact nearby genes

After determining that pAB3 integration does not balance chromosomes, it was next hypothesised that pAB3 integration alters the transcription of nearby genes and/or *tetR1*/*tetR2*. Therefore, RNAseq was performed on non-drug and meropenem-exposed mff and AB042 isolates. AB042 was selected as it was representative of other ST437 GS2.24 isolates. We found no clusters of DEGs near the pAB3 boundaries, and only *tetR2* was found to be very slightly statistically significantly upregulated under meropenem exposure (Fig 4).

**Fig 4.**
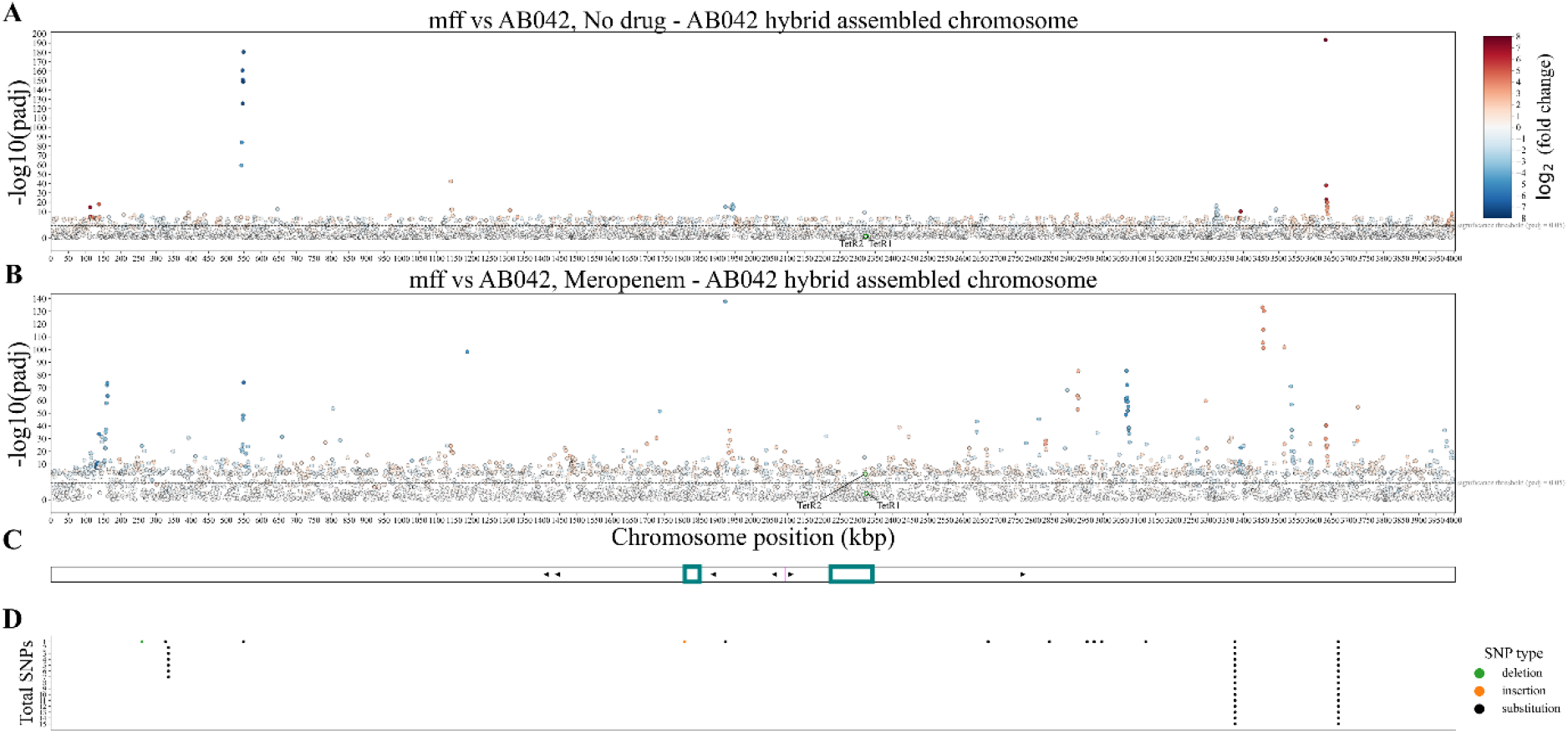
DEGs between mff and AB042. (A) and (B) are Manhattan plots with each point representing a gene and the dashed line showing the significance threshold. DEGs between mff and AB042 were determined separately for untreated (A) and meropenem-exposed (B) isolates, with three biological replicates per strain and condition. (C) shows a map of the hybrid assembled AB042 chromosome. rRNA operons are represented by black arrows. *dnaA* is shown in magenta. pAB3 sequences are shown with teal rectangles. (D) shows SNP differences between mff and AB042 chromosomes coloured by type as indicated by the legend.

Numerous DEGs were identified in the non-drug condition (Fig 4A), mostly related to the Csu pili and lipid metabolism. However, it should be noted that AB042 is known to have a deletion in the *adeN* gene, which encodes a repressor of the *adeIJK* operon (43,44). This deletion likely triggers AdeIJK overexpression, which is known to cause Csu pili downregulation (45) and altered lipid metabolism (46). Therefore, these DEGs are likely the result of *adeN* perturbation rather than pAB3 integration.

### pAB3 contains the ARG *sul2* which is maintained on the AB042 and VU chromosome

After finding that some chromosomes had lost portions of the integrated pAB3 sequence, it was noted that a portion of pAB3 sequence was retained on the chromosome. The only ARG detected on the hybrid mff pAB3 assembly was *sul2* in line with previous literature (3). The mff chromosome encoded no *sul2*, while VU and AB042 chromosomes encode a single copy located in the 50 kbp pAB3 block. Therefore, it seems probable that VU has maintained the sulfamethoxazole-resistant phenotype conferred by pAB3 while also potentially having a T6SS+ phenotype due to the loss of the 100 kbp pAB3 block containing *tetR1* and *tetR2*.

## Discussion

ATCC 17978 is one of the most commonly used strains for *A. baumannii* research, and the previously observed integration of the large conjugative plasmid pAB3 into the chromosome was widely considered to result from an assembly error in the original genome (3,5–7,15,16). Here, we show that pAB3 does integrate into the ATCC 17978 chromosome, and is found as two blocks of 50 kbp and 100 kbp in length. This integration is also associated with a large inversion approximately symmetrical around the oriC, with IS*Aba18* acting as the IR, resulting in a structural arrangement classified as GS2.24 by Socru.

To confirm that the integration observed was not the result of assembly errors, three isolates (mff, AB042, and VU) were whole-genome sequenced with both long-read and short-read technologies, then hybrid assembled. In AB042, integration of pAB3 and the GS2.24 structural arrangement was observed as in the publicly available chromosomes. In VU, the GS2.24 structural arrangement and a 50 kbp pAB3 block was observed, but the 100 kbp block (containing *tetR1* and *tetR2*) was missing. Notably, we additionally found two instances in publicly available genomes where the 100 kbp block had lost *tetR2*. We speculate that the loss of some portion of the integrated pAB3 sequence may be the next stage in the structural variation of the ATCC 17978 chromosome, with *tetR2* (and potentially *tetR1*) lost in order to de-repress the T6SS.

Based on the results presented above, the sequence of genome rearrangements occurring in the ATCC 17978 chromosome can be reconstructed and is summarised in Fig 5. Firstly, an ATCC 17978 chromosome must acquire at least one chromosomal copy of IS*Aba18*. This is observed in the mff strain, but not in other ST437 GS1.0 genomes (Fig 5A). Second, the large conjugative plasmid pAB3 integrates into the chromosome (Fig 5B), likely involving the segments which derived from a common ancestor and therefore containing high sequence similarity (5). Thirdly, the integrated pAB3 supplies a second IS*Aba18*, which can then act as an IR mediating an inversion resulting in the GS2.24 structural arrangement (Fig 5C). This splits the pAB3 sequence into 100 kbp and 50 kbp blocks. A fourth putative stage is loss of some or all of the 100 kbp block containing *tetR2* (and potentially *tetR1*), which in turn de-represses the T6SS. This stage leaves behind a portion of the pAB3 sequence encoding *sul2* which confers a sulfamethoxazole-resistant phenotype (Fig 5D).

**Fig 5.**
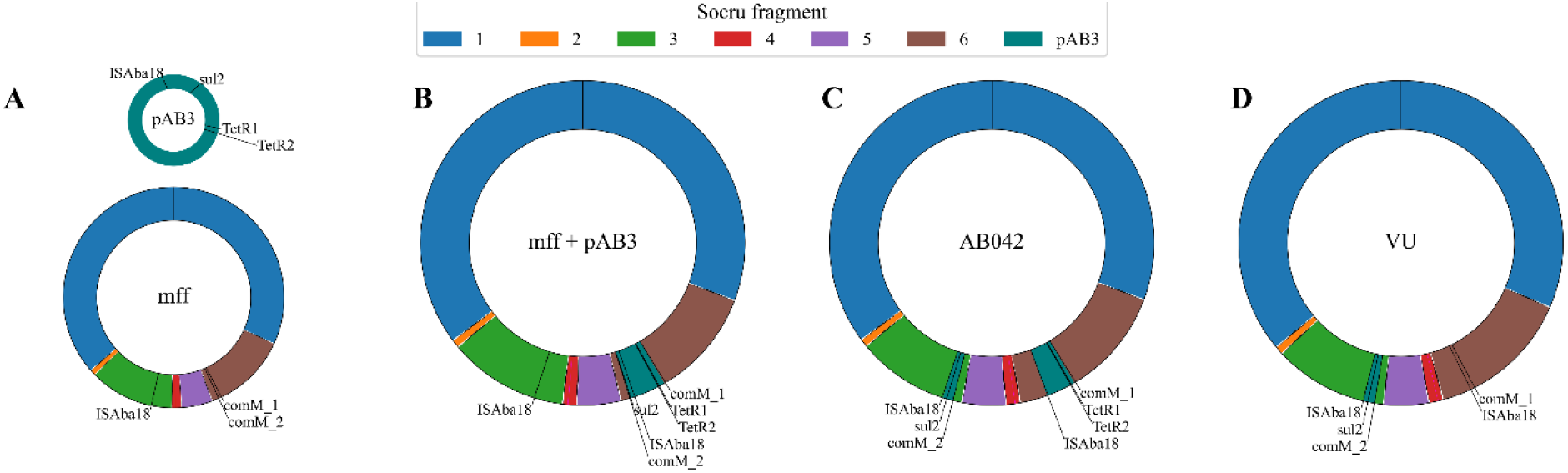
Structural variations in the ATCC 17978 chromosome. (A-D) represent sequential stages in chromosome structure as described in the text. Socru fragments are colour coded as per the legend and rRNA operons are shown with black arrows indicating the direction determined by Socru. pAB3 sequences are shown with teal boxes. The *dnaA* gene, located in fragment 4, is shown in magenta.

It was initially hypothesised that the integration of pAB3 and subsequent inversion may act to balance the chromosome. However, the opposite was found to be the case. Nonetheless, we found that in all ST437 chromosomes the *dnaA* gene was over 45% of the chromosome length from the ter, well within the range of non-ST437 chromosomes and unlikely to result in fitness loss based on previous literature (12,13). For instance, in *E. coli* an imbalance of 5-10% (oriC 45-40% from the ter) has no effect on growth or colony morphology (12), and in *S. Typhi* an imbalance of 198.5° (oriC ∼45% from ter) caused no decrease in growth rate compared to a wild type (WT) strain with an imbalance of 177.6° (oriC ∼49% from ter) (13). Therefore, it is likely that this imbalance results in no fitness loss.

It was also hypothesised that integration of pAB3 may impact the transcription of nearby genes and/or impact the baseline expression of *tetR1* and *tetR2*. However, we did not find this to be the case, and only *tetR2* was differentially expressed between mff and AB042 under meropenem exposure. This could indicate that pAB3 integration alters the regulation of *tetR2* to increase its upregulation under antibiotic exposure, thereby improving the repression of the T6SS. However, AB042 has been documented to have an *adeN* deletion (43,44), and DEGs identified in the non-drug condition were attributable to this deletion. Therefore, further analysis of DEGs between isolates is confounded by the *adeN* deletion in AB042.

Based on the transcriptional data and the observed loss of *tetR2*-containing sections in several chromosomes, we speculate that pAB3 integration may make the deletion of the *tetR2* gene (and in some cases the *tetR1* gene) more tractable. We also note that this process maintained the only ARG detected on pAB3, *sul2*, in VU and in publicly available genomes this involves the loss of only a small portion of pAB3, leaving the rest in the chromosome. Therefore, we speculate that these deletions enable the activation of the T6SS while maintaining a sulfamethoxazole-resistant phenotype, which would not be possible if the entirety of pAB3 was lost. The current model is that ATCC 17978 can have only T6SS+ or sulfamethoxazole-resistant phenotypes depending on pAB3 presence/absence (3). However, the integration of pAB3 may enable further rearrangements which allow for a T6SS+ sulfamethoxazole-resistant phenotype. This mechanism also enables the retention of additional genes encoded on pAB3 which may confer additional fitness benefits beyond sulfamethoxazole resistance.

In the dataset analysed here it was found that just over half of the ST437 chromosomes had both pAB3 integration and a large inversion, both of which had not been previously identified. This is despite the ATCC 17978 strain being extensively used in laboratories and sequenced many times. This can be attributed to some isolates genuinely lacking pAB3 or having it present as a separate plasmid, and the focus on small-scale SNPs when analysing genomic differences between isolates. This highlights the need for further attention to large-scale SVs, which may be far more prevalent than previously thought, even in well studied strains. It should also be noted that the loss of the 100 kbp block in VU seems to have occurred during transport between laboratories, similar to previous reports on the loss of pAB3 (6), showing large-scale SVs can occur rapidly. Finally, this shows that the ATCC 17978 genome is much more varied than previously known, adding to the concerns around the widespread use of this strain in laboratories without specific confirmation of the genotype (7).

In conclusion, we have used a combination of publicly available and hybrid assembled ST437 genomes to confirm that the large conjugative plasmid pAB3 integrates into the ATCC 17978 chromosome. This integration enables a large inversion by providing a second copy of IS*Aba18* which acts as an IR, resulting in the chromosomal arrangement labelled GS2.24 by Socru. No evidence was found that pAB3 integration alters the baseline transcription of nearby genes or the *tetR1*/*tetR2* genes. However, several instances were found where portions containing *tetR2* of the integrated pAB3 sequence were lost, suggesting integration may make further rearrangements more tractable, enabling a putative T6SS+ sulfamethoxazole-resistant phenotype.

## Author Contributions

MN: conceptualisation, data curation, formal analysis, investigation, methodology, visualisation, writing – original draft; GCL conceptualisation, methodology, supervision, writing – review and editing; BAE: conceptualisation, funding acquisition, investigation, methodology, project administration, resources, supervision, writing – original draft, writing – review and editing.

## Acknowledgments

We would like to thank Mario Feldman and Ayush Kumar for providing strains used in this study. We would like to thank Sadaf Farooq for MIC testing of strains. Thank you to Claire Hill for useful comments on genomic rearrangements.

